# Spatial distribution of epibionts on olive ridley sea turtles at Playa Ostional, Costa Rica

**DOI:** 10.1101/669523

**Authors:** Nathan J. Robinson, Emily Lazo-Wasem, Brett O. Butler, Eric A. Lazo-Wasem, John D. Zardus, Theodora Pinou

## Abstract

There is a wealth of published information on the epibiont communities of sea turtles, yet many of these studies have exclusively sampled epibionts found only on the carapace. Considering that epibionts may be found on almost all body-surfaces and that it is highly plausible to expect different regions of the body to host distinct epibiont taxa, there is a need for quantitative comparative studies to investigate spatial variation in the epibiont communities of turtles. To achieve this, we measured how total epibiont abundance and biomass on olive ridley turtles *Lepidochelys olivacea* varies among four body-areas of the hosts (n = 30). We show that epibiont loads on olive ridleys are higher, both in terms of number and biomass, on the skin than they are on the carapace or plastron. This contrasts with previous findings for other hard-shelled sea turtles, where epibionts are usually more abundant on the carapace. Moreover, the arguably most ubiquitous epibiont taxon for other hard-shelled sea turtles, the barnacle *Chelonibia spp*., only occurs in relatively low numbers on olive ridleys, while the barnacles *Stomatolepas elegans* and *Platylepas hexastylos* are far more abundant. We postulate that these differences between the epibiont communities of different sea turtle taxa could indicate that the carapaces of olive ridley turtles provide a more challenging substratum for epibionts than do the hard shells of other sea turtles. In addition, we conclude that it is important to conduct full body surveys when attempting to produce a holistic qualitative or quantitative characterization of the epibiont communities of sea turtles.

## INTRODUCTION

In the marine environment, almost any non-toxic, non-protected surface will eventually be colonized by an array of microorganisms, plants, algae, or animals [1,2]. This is true for non-living substrata, such as rocks, sand grains, or the shells of dead molluscs, as well as the bodies of living marine animals. Those organisms that live on the surfaces of other organisms are referred to as epibionts [3], a grouping of diverse taxa with a wide-range of life-histories, physiological tolerances, and mechanisms for attaching to their host [4]. It is therefore not surprising to discover that epibiont communities vary among different host species [5,6], populations [7], and even between different body-areas on a single host [8,9]. Understanding the factors driving spatial variation in the distribution of epibionts, be it on the scale of an ocean or a host’s body, can shed light the habitat requirements of these varied associates. In turn, this information can be used to understand habitat preferences or behaviour of the host [e.g. 10,11] and even epibiont-related disease transmission [e.g. 12].

Sea turtles arguably host the most abundant and diverse epibiont communities of all large marine megafauna. In an effort to characterize these epibiont communities, researchers have generated hundreds of species-lists reporting the presence of epibiont taxa for sea turtle populations around the world [13]. As these datasets have grown, there is now an impetus to synthesise these data and elucidate epibiont community spatial patterns globally and make quantitative assessments of epibiont loads [14]. Yet to help draw robust conclusions about geographic differences in epibiont communities, it would be beneficial to understand spatial patterns in epibiosis at a finer-scale – the scale of a turtle’s body. Indeed, it is highly likely that epibionts are non-uniformly distributed over a turtle host for several reasons. For example, the physical properties of certain body-parts, such as the carapace, differ from those of the plastron or skin [15], which likely affects attachment or settlement of different epibiont species. Different locations on the host’s body may also present different opportunity for feeding. For example, some epibiont species feed preferentially on certain tissues and thus will be found predominantly in those areas [16] whereas filter feeders might prefer to be in areas where water flow over the host is optimal [17]. Either of these factors, as well as attachment mode, crypsis, mating habits, and others, could lead to distinct variation in the distribution of epibionts on sea turtles.

While it may be clear that epibiont distribution is likely to vary over the host’s body, many studies on sea turtle epibiosis have either not provided any quantitative assessment of where on the host the epibionts were found [e.g. 7,18], and/or have simply sampled epibionts from a single location on the animal, commonly the carapace [e.g. 19,20]. Tellingly, even some of the most seminal studies investigating the distribution patterns of epibionts on the loggerhead turtle *Caretta caretta* and green turtle *Chelonia mydas* hosts have focused exclusively on the carapace [8,21]. There are only two studies of which the authors are aware that have compared the abundance of epibionts from over the entire body: Devin and Sadeghi (2010) [22] and Razaghian et al. (2019) [23]. Interestingly, both studies, which focused on hawksbill turtles *Eretmochelys imbricata* nesting in Iran, identified that epibiotic barnacles were actually more common on the plastron than the carapace. If this is true for all sea turtle species, then this finding could have important implications for epibiont sampling. Specifically, if epibionts are not uniformly distributed on the hosts’ body then the actual biodiversity of epibionts might be underestimated if only sampled from a single region of the body.

To better understand spatial variation in epibiont communities on sea turtle hosts, we made a quantitative assessment of the abundance and biomass of several epibiotic taxa from different regions of the body of olive ridley sea turtles *Lepidochelys olivacea*. This species was chosen because several studies have focused on characterizing the epibiont communities of this species in recent years [e.g. 6,11,18,24] yet no previous studies have specifically looked at epibiont distributions for this host species. Our null prediction was that there would be no variation in the abundance or total biomass of the various epibiont taxa among the different body-areas.

## METHODS

### Study Site

Olive ridley turtles were sampled from Playa Ostional, which is located on the Pacific coast of the Nicoya Peninsula in northwest Costa Rica (9° 59’ N, 85° 42’ W). We selected this site as it is one of a few beaches worldwide that has mass-nesting assemblages or “*arribadas*” of olive ridley turtles, thus making it feasible to sample large numbers of turtles relatively easily. These arribadas generally occur once a month and last three to eight days [25]. At Ostional, over 500,000 turtles have been estimated to nest in a single arribada event [26] at densities of over 4 clutches m^-2^ [27]. For this study, we sampled epibionts from a total of 30 turtles spanning three separate arribada events. Specifically, we sampled 10 turtles on 6-Dec-2015, 5 turtles on 1-Jan-2016, and 15 turtles on 7-Feburary-2016.

### Sampling Protocol

During arribada events, we patrolled the beaches of Ostional to encounter nesting turtles. To avoid any possibility that sampling would interrupt the nesting process, we only examined turtles after they had completed oviposition. In addition, we sampled only animals that appeared to be in good health with no visible injuries, lesions, or tumours as certain epibionts are often found in association with certain injuries [28,29].

When a suitable turtle was encountered, we measured its Curved Carapace Length using a flexible tape measure. The turtle’s flippers were then restrained by hand and the animal was moved onto a plastic tarp. This limited the ability of the turtle to flick sand on its carapace, which would impede efforts to sample epibionts. We then exhaustively collected epibionts by scraping or prying them off the turtle using a knife or tweezers. After all visible epibionts were collected from the dorsal surfaces of the turtle, we would briefly (< 10 mins) flip the animal onto its carapace to collect epibionts from its ventral surfaces. All epibionts were divided between separate containers by the region of the body where they were collected. We divided the body into four regions: (1) the head, shoulder, and fore flippers (subsequently termed head), (2) the tail, cloaca, hind flippers, and inguinal cavities (subsequently termed tail), (3) the carapace, or (4) the plastron (Figure 1). Epibiont samples were preserved in 75% non-denatured ethanol following the protocols outlined in Lazo-Wasem et al. (2011) [18]. No attempt was made to quantify the abundance or distribution of micro-epibionts, such as diatoms, even though it is known that they exist in high abundances on sea turtles [6,24]. We did not tag any of the turtles, yet due to high numbers of individuals at each arribada it was easy to ensure that the same individual was not sampled repeatedly.

**Figure 1.**
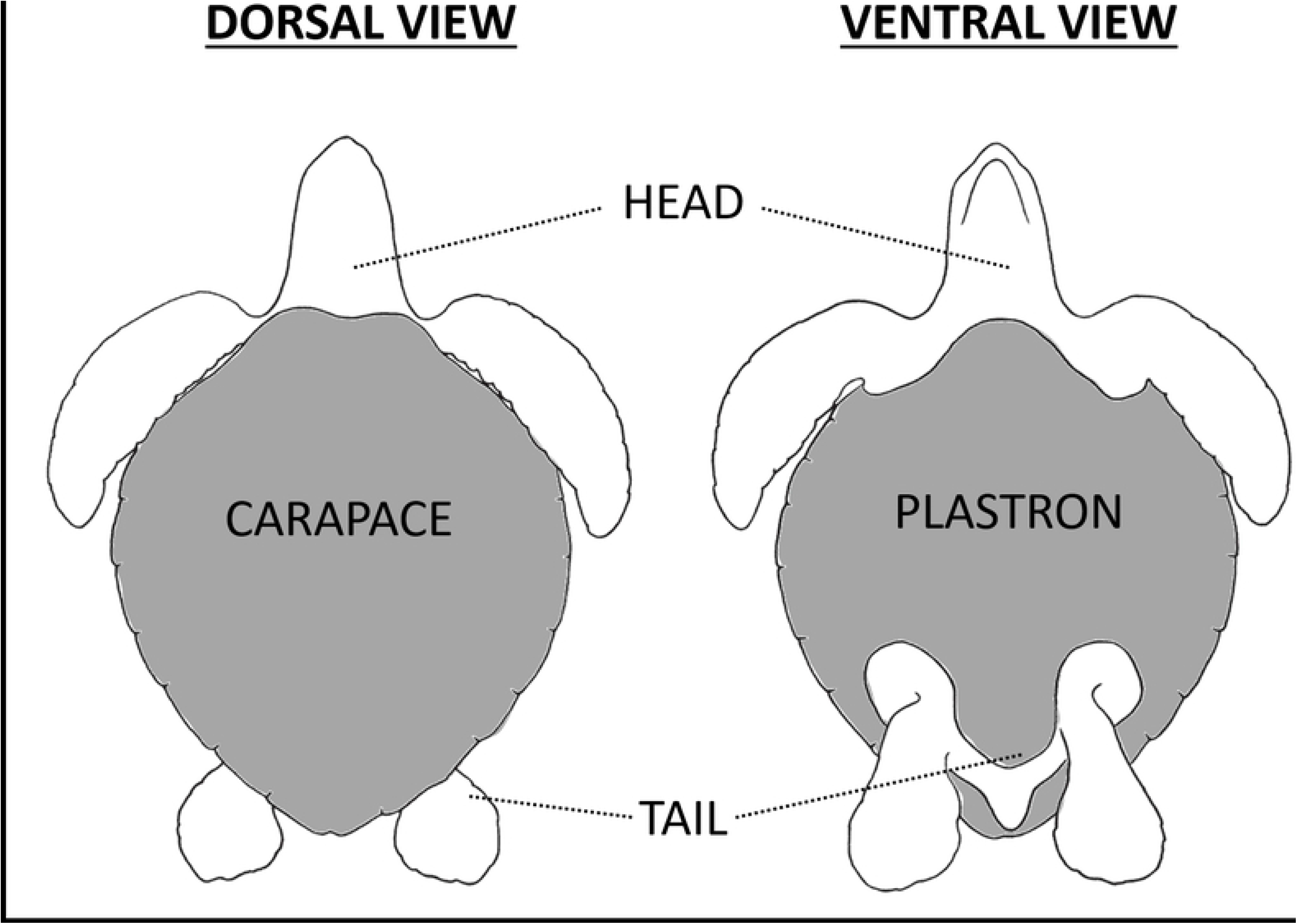
An illustration of the four different regions of the body that were sampled for epibionts. (1) The head, shoulder, and fore flippers (= **head**), (2) The tail, cloaca, hind flippers, and inguinal cavities (= **tail**), (3) The shell dorsum (= **carapace**), and (4) The shell ventrum (= **plastron**),.

All epibionts were identified to the lowest taxonomic level by consulting appropriate literature. The samples were then enumerated, dried, and weighed to the nearest 0.00001g using a Mettler B6 Semi-Micro Balance. Samples were catalogued and deposited in the Yale Peabody Museum of Natural History, U.S.A. Specimen records are available at http://peabody.yale.edu/collections.

To test our null prediction that there would be no variation in the abundance or total biomass of the epibiont taxa between the different regions of the turtles’ body, we used Kruskal-Wallace tests with ad-hoc pair-wise Mann-Whitney tests. Separate tests were run for each major epibiont taxa. Kruskal-Wallace and Mann-Whitney tests were conducted using PAST V.3.13, confirming significance when p ≤ 0.05. We chose these non-parametric tests, over their parametric counterparts, because the data included many zeros and were not normally distributed.

## RESULTS

Of the 30 olive ridley turtles that were sampled (CCL: 66 – 69 cm range), epibionts were found on all but one individual. In total, we collected 1614 individual epibionts, which collectively weighed 14.01 g. The mean number of epibionts per host was 53.80 with a mean cumulative mass of 0.47 g. The epibiont load by body region (considering all epibiont taxa) was highest on the tail with 59.23 % of epibionts being found in this location and constituting 64.11 % of the total epibiont biomass (Figure 2). The location with the next highest epibiont load was the head with 31.69 % of epibionts being found in this location and constituting 24.83 % of the total epibiont biomass. For both the carapace and plastron, epibiont load was very low and never exceeded more than 8 % of the total epibiont load.

**Figure 2.**
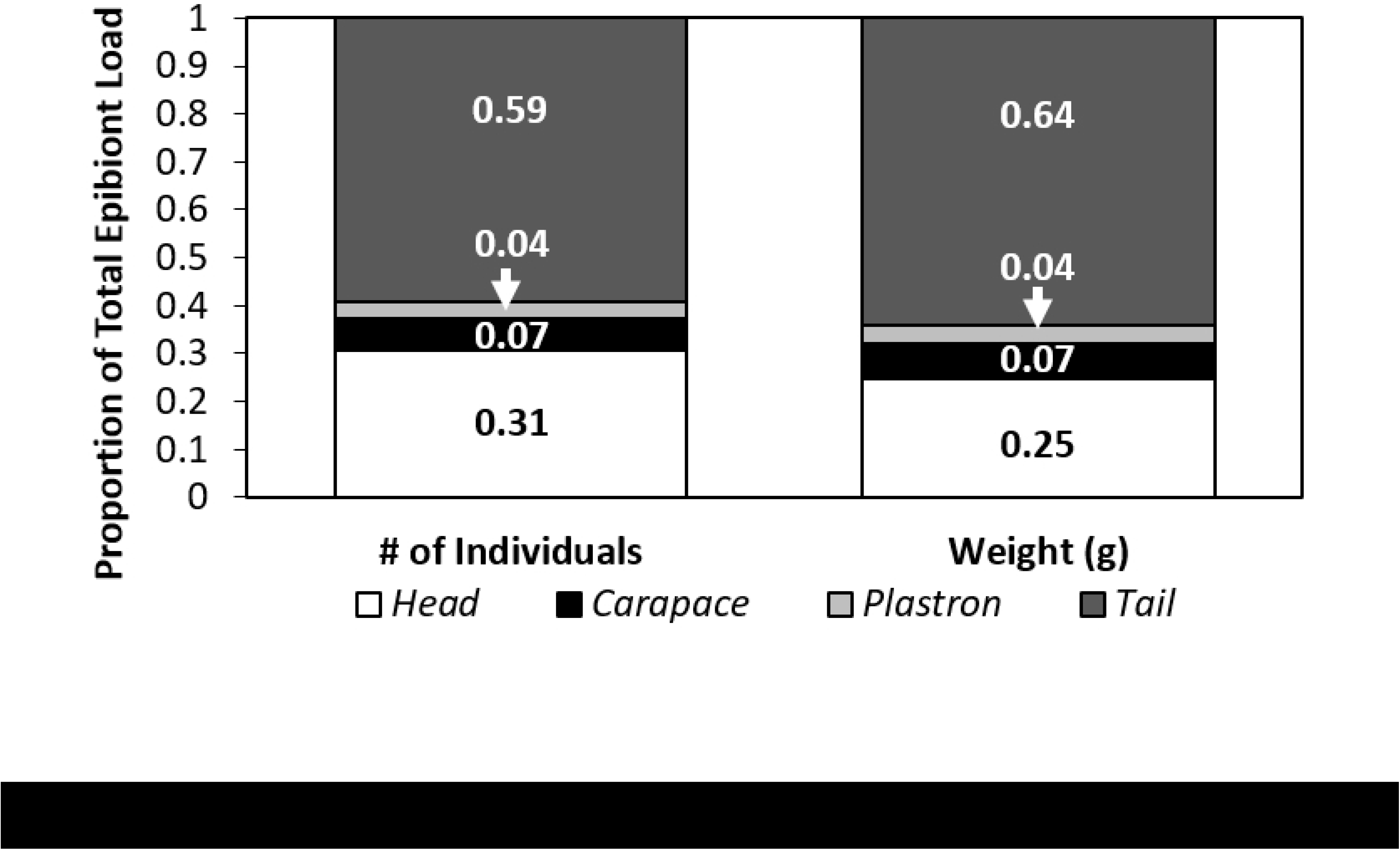
Proportion of the total epibiont load found on the four body regions (head, carapace, plastron, and tail) of olive ridley sea turtles. The left graph represents % abundance and the right graph represents % total biomass.

We conservatively estimate that the diversity of epibionts represented 20 different taxa (see Table 1 for a full list). Most of these taxa were relatively rare and only five were found in mean abundances exceeding one individual per host. In alphabetical order these were: *Balaenophilus manatorum*, *Chelonibia testudinaria*, *Platylepas hexastylos*, *Stomatolepas elegans*, and small individuals of family Platylepadidae that cannot as yet be identified. This platylepadid may comprise more than one species, but for now the most conservative approach is to view it as a single operational taxonomic unit.

**Table 1.** Mean # and biomass of the different epibiont species found on 30 olive ridley sea turtles sampled from Playa Ostional, Costa Rica.

| Systematic Group | Epibiont Taxon | Mean # of epibionts per host | Mean epibiont biomass per host (g) |
| --- | --- | --- | --- |
| Annelida: Hirudinea | *Ozobranchus branchiatus* | 0.1 | < 0.1 |
| Annelida: Polychaeta | Polychaeta | < 0.1 | < 0.1 |
| Annelida: Polychaeta | Serpulidae | 0.1 | < 0.1 |
| Chlorophyta | Algae / Hydroid | 0.3 | 0.1 |
| Chelicerata: Pycnogonida | Pycnogonida | < 0.1 | < 0.1 |
| Crustacea: Amphipoda | Caprellidea | 0.1 | < 0.1 |
| Crustacea: Amphipoda | Corophiidae | < 0.1 | < 0.1 |
| Crustacea: Amphipoda | *Podocerus chelonophilus* | < 0.1 | < 0.1 |
| Crustacea: Brachyura | *Planes* spp. | < 0.1 | < 0.1 |
| Crustacea: Cirripedia | *Chelonibia testudinaria* | 2.8 | 2.3 |
| Crustacea: Cirripedia | Chelonibiidae | < 0.1 | < 0.1 |
| Crustacea: Cirripedia | *Conchoderma virgatum* | < 0.1 | < 0.1 |
| Crustacea: Isopoda | Corallanidae | < 0.1 | 0.1 |
| Crustacea: Cirripedia | *Balanoidea* sp. | 0.5 | < 0.1 |
| Crustacea: Cirripedia | Platylepadidae spp. | 2.1 | 0.1 |
| Crustacea: Cirripedia | *Platylepas hexastylus* | 15.7 | 2.4 |
| Crustacea: Cirripedia | *Platylepas* sp. | 0.2 | < 0.1 |
| Crustacea: Cirripedia | *Stomatolepas elegans* | 20.9 | 8.9 |
| Crustacea: Copepoda | *Balaenophilus manatorum* | 8.7 | < 0.1 |
| Crustacea: Copepoda | Harpacticoida | < 0.1 | < 0.1 |
| Mollusca: Gastropoda | Unknown sp. | < 0.1 | < 0.1 |

The most common taxon was *Stomatolepas elegans*, which constituted 38.85 % and 63.30 % of the total epibiont load in terms of abundance and mass respectively. *Platylepas hexastylos* was almost as abundant, constituting 29.30 % of all epibionts but individuals were generally lower in mass than *Stomatolepas elegans* and so constituted only 17.24 % of the total epibiont biomass. In contrast, *Chelonibia testudinaria* was not very abundant, comprising only 5.27 % of all epibionts, yet it constituted 16.54 % of total epibiont biomass. *Balaenophilus manatorum* was relatively abundant, constituting 17.66 % of all epibiont individuals, yet the combined biomass of this species was less than 0.01 g and so relatively negligible. Similarly, *Platylepadidae* spp. *c*onstituted 5.70 % of the total epibiont abundance yet only 0.8 % of the total epibiont biomass, further indicating that these individuals were very small. The remaining 15 taxa constituted only 3.34 % and 2.09 % of the total epibiont load in terms of abundance and biomass respectively (Figure 3).

**Figure 3.**
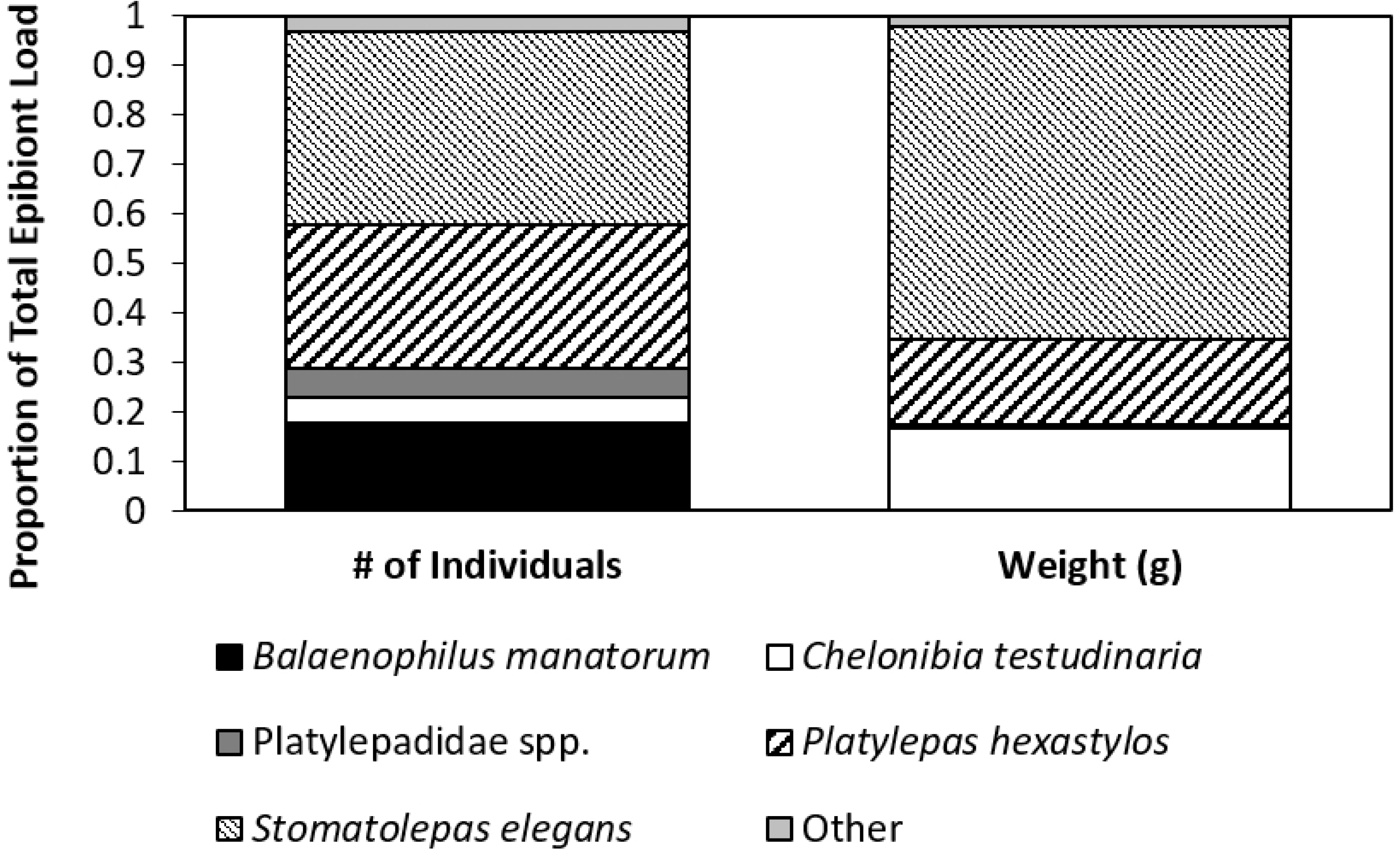
Proportion of the five main epibiont species relative to the total epibiont load found on the entire body of olive ridley sea turtles. The left graph represents % abundance and the right graph represents % total biomass.

Kruskal-Wallis Test tests indicated significant differences between both epibiont abundance and biomass for the different body regions (abundance: H_(χ2)_ = 23.42, d.f. = 3, p < 0.0001; biomass = H_(χ2)_ = 16.42, d.f. = 3, p < 0.0001). Thus, we can reject the null prediction that epibionts are uniformly distributed on their hosts (Figure 4).

**Figure 4.**
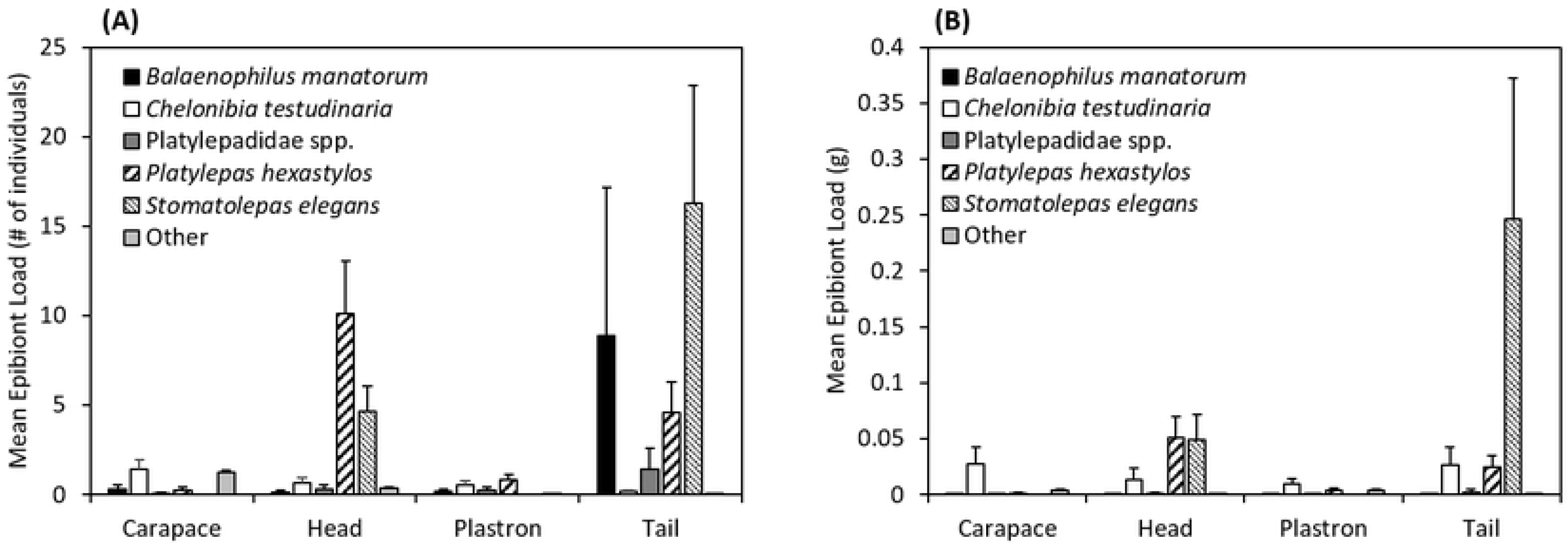
Mean number of individuals (A) and mean biomass (B) for the five main epibiont species (*Balaenophilus manatorum*, *Chelonibia testudinaria*, Platylepadidae spp., *Platylepas hexastylos*, and *Stomatolepas elegans*) from the four 4 different body regions (head, carapace, plastron, and tail) of olive ridley sea turtles.

When we tested for differences in the abundance or biomass of the five major epibiont taxa between the different body region, we observed significant differences in the abundance and biomass of *Stomatolepas elegans* (abundance: H_(χ2)_ = 25.49, d.f. = 3, p < 0.0001; biomass = H_(χ2)_ = 15.67, d.f. = 3, p < 0.0001) and *Platylepas hexastylos* (abundance: H_(χ2)_ = 16.87, d.f. = 3, p < 0.0001; biomass = H_(χ2)_ = 15.67, d.f. = 3, p < 0.0001). Ad-hoc pair-wise Mann-Whitney tests revealed that for both *Stomatolepas elegans* and *Platylepas hexastylos* there were no statistical differences in either abundance or biomass between head or tail sections (p < 0.0001 for all tests). Instead, statistical differences were observed in abundance or biomass between both the head and tail when compared to either carapace or plastron (p > 0.5 for all tests).

No significant differences were observed in the abundance or biomass of *Chelonibia testudinaria* (abundance: H_(χ2)_ = 2.94, d.f. = 3, p = 0.1973; biomass = H_(χ2)_ = 1.613, d.f. = 3, p = 0.4650)*, Balaenophilus manatorum* (abundance: H_(χ2)_ = 0.2748, d.f. = 3, p = 0.5139; biomass = H_(χ2)_ = 0.2727, d.f. = 3, p = 0.5173), and Platylepadidae spp. (abundance: H_(χ2)_ = 0.4883, d.f. = 3, p = 0.5830; biomass = H_(χ2)_ = 0.4834, d.f. = 3, p = 0.5873) between the different body areas.

## DISCUSSION

### Spatial distribution of epibionts

Knowledge of how sea turtle epibionts are distributed on their hosts can provide valuable insights into the ecology and habitat requirements of these extracorporeal companions. Furthermore, when epibionts are not uniformly distributed over the host then sampling only from certain body-parts runs the risk of under-sampling and misrepresenting certain epibiont taxa in qualitative assessments and certainly in quantitative evaluations. To test this concern, we measured how the abundance and biomass of epibionts varies between different body regions on host olive ridley turtles. We discovered that epibiont loads on olive ridley turtle are highest, both in terms of number and biomass, on the soft tissues of the head and tail than they are on the carapace or plastron. Specifically, 60 % of the total number of epibionts was found on the tail and 32 % was found on the head. In contrast, only 6 % of the total number of epibionts was found on the carapace and 4 % on the plastron. This finding suggests that studies exclusively sampling epibionts from the carapace, or any other specific, body-part could significantly under sample the total epibiont load and run the risk of not detecting or at least underrepresenting those taxa that are not commonly found on the sampled body-part. For example, the most common epibiont taxon recorded in this study, *Stomatolepas elegans* was absent from the carapace or plastron even though found in high abundance on the tail and head. Somewhat predictably, due to the intimate association between *Balaenophilus manatorum* and *Stomatolepas elagans* [28], *Balaenophilus manatorum* was also only reported in substantial numbers on the tail section where *Stomatolepas elagans* was equally common and barely recorded elsewhere on the turtles. Similarly, *Platylepas hexastylos* was over 5 times more abundant on the head and tail than it was on the carapace or plastron.

It is also interesting to note that the total biomass of *Stomatolepas elegans* and *Platylepas hexastylos* individuals exceeded that of *Chelonibia testudinaria* despite individuals of the latter being heavier. Indeed, individual *Stomatolepas elegans* and *Platylepas hexastylos* weighed an average of 0. 4 g and 0.2 g respectively, while *Chelonibia testudinaria* weighed on average 0.8 g. Thus, from an ecological standpoint the small, less conspicuous epibionts, scattered in high densities in the soft-tissues, and consequently more difficult to find, may be the most ecologically important for the host animal.

Thus, it is clear that to get a true understanding of the epibionts on sea turtles, studies should sample all body-surfaces. This is necessary to build both an understanding of the habitat requirements of epibiont species and a methodology for using them as biological indicators. For example, *Balaenophilus manatorum* has been previously suggested as a vector for fibropapillomatosis [28] and this prediction has been further supported by more recent studies [16,30]. The presence of this species could therefore be important to monitor in reference to turtle health. Yet if researchers only sample epibionts from the carapace and plastron, they limit their capacity to identify the presence or absence of this species. We therefore recommend that future studies attempt to survey all body-sections on turtles when either attempting to characterize the entire epibiont community quantitatively or when specifically trying to determine the presence or absence of taxa qualitatively.

### Differences between host species

The discovery that epibiont loads are higher for olive ridley turtles on the soft tissues, such as the head and tail, relative to the carapace and plastron was somewhat unexpected. Indeed, comparable studies that conducted quantitative assessments of epibiont distributions on hawksbill turtles found the majority of epibiotic barnacles were found on the plastron and carapace respectively [22,23]. Furthermore, it has often been assumed although there is little quantitative data to support it, that for loggerhead and green turtles the highest quantities of epibionts are found on the carapace [see 31].

The distinct difference in the distribution of epibionts between these different sea turtle species begs the question of why such inter-specific differences should occur. One hypothesis could be that these discrepancies are driven by differences in the behaviour of each host species, exposing them to settlement by different larval epibionts with different habitat preferences (i.e. reef, seagrass meadow, or soft sediment), yet we consider this to be unlikely considering the large range and behaviour overlaps between many turtle species. Instead, we propose that differences in the physical properties (e.g. microstructure, chemical composition, physical properties) of the carapace and dermis among different turtle species drive epibiont settlement.

Supporting the notion that olive ridley carapaces are a challenging substrate for epibionts settlement is our observation of the relative paucity of *Chelonibia* barnacles associated with this host*. Chelonibia* spp. are common taxa on the carapaces of loggerhead turtles [19, 20], hawksbill turtles *Eretmochelys imbricata* [32], and green turtles [6]; however, we observed a mean of only 2.8 individuals of *Chelonibia testudinaria* per host, which is far lower than the observed 16.8 individuals per host on green turtles that share nesting beaches with olive ridley turtles in Costa Rica [6]. The *Chelonibia testudinaria* found on olive ridley turtles appeared far smaller than those on these green turtles (NJ Robinson, personal observation) suggesting that maybe they are not surviving long enough on this host to grow to their full size. In contrast, the most common epibiont observed on olive ridley turtles was *Stomatolepas elegans*, which unlike *Chelonibia testudinaria*, tends to bury into the softer tissues using an array of plates to hold itself in place.

### Geographic patterns in epibiont communities in olive ridleys

In recent years, the epibiont communities of olive ridley sea turtles in the East Pacific Ocean have been described in several studies [6,11,18,24]. Collectively between these four studies, a total of 291 olive ridley turtles have been sampled for epibionts. From this extensive dataset, it is clear that while the epibiont communities of olive ridley turtles in the East Pacific are diverse and new epibiont associations have been revealed in each study, these epibiont communities are generally dominated by a few main taxa. The taxa include: the barnacles *Chelonibia testudinaria*, *Conchoderma virgatum*, *Lepas* spp. *Platylepidaeas decorata* (which we describe as Platylepadidae spp. in study for reasons described earlier), *Platylepas hexastylos*, and *Stomatolepas elegans*; the leech *Ozobranchus branchiatus*, the amphipod *Podocerus chelonophilus*, and the copepod *Balaenophilus manatorum*. All other taxa tend to be recorded at relatively low frequencies (i.e. less than one per 50 hosts).

This is not to say, however, that these more commonly observed taxa for East Pacific olive ridley turtle are found on every turtle. In fact, even these commonly observed epibiont taxa are noticeably absent from some studies. For example, *Podocerus chelonophilus* was observed on more than 25 % of the olive ridley turtles sampled both on the beach of Playa Grande [6] and in the nearby waters El Coco [11]. In this study on Playa Ostional however, we recorded a single individual only of *Podocerus chelonophilus* among all 30 turtles, even though this beach is less than 50 km away from the previously mentioned sampling locations. Majewska et al. (2015) also reported a similarly low prevalence of *Podocerus chelonophilus* from olive ridley turtles sampled at Playa Ostional, with only 4 % of all the turtles hosting this epibiont [24]. *Lepas* spp. are also noticeably less prevalent at Playa Ostional, relative to other nearby locations [6,11,24].

One possible hypothesis to explain differences in the abundances of these common epibiont taxa between turtles sampled on Playa Ostional compared to those sampled at other locations nearby is that Playa Ostional experiences different oceanographic conditions. While previous studies have indeed noted the strong presence of offshore currents on Playa Ostional, relative to other nesting beaches on the Pacific coast of Costa Rica [33], it seems unlikely that this is the primary factor driving the differences in epibiont communities between these locations. This is because nesting olive ridley turtles are known to move over larger distances between nesting events than the distances between these beaches (< 50 km) [34], and thus there is likely significant overlap in the inter-nesting habitats of these turtles even though they nest on different beaches. Instead, we propose that a more robust hypothesis is that the differential prevalence of certain epibiont taxa is linked to differences in behaviour between turtles nesting at each site. Indeed, Playa Ostional is a well-known arribada site where thousands of turtles aggregate in mass nesting events, while Playa Grande and the waters of El Coco likely host turtles that are solitary nesters. Perhaps during such mass mating events there is a much greater chance for epibionts to be scraped off than there is in smaller mating events, meaning the arribada nesting turtles have fewer epibionts. Alternatively, as many epibionts, such as *Podocerus chelonophilus* and *Balaenophilus manatorum* are not free swimming and must likely be transferred on contact between hosts [30], it could be that the great potential for the transfer of individuals between hosts means that they even though they might be found on more hosts, they occur in lower numbers per host. Understanding the factors that are driving distinct variation in the presence and absence of these common epibiont species would be an interesting avenue for future research and could reveal several aspects about the behaviour of their hosts that might be challenging to detect using more conventional methods [30,35].

## ACKNOWLEDGEMENTS

Epibiont collection was conducted under MINAE permits (ACT-ORDR-099-15 and ACG-PI-058-2015). This research was performed in accordance with the Purdue University Animal Care and Use Committee. Samples were exported from Costa Rica under a SINAC permit (DGVS-183-2016). Pilar Santidrián Tomillo, Christian Diaz Chuquisengo, Elizabeth Solano, and Myriam Norori helped with permitting. Lourdes Rojas aided with identifying taxa.

## AUTHOR CONTRIBUTIONS

NJR wrote the paper with input from all authors. Conceived and designed the experiments: NJR, EAL-W, JDZ, and TP. Collected samples in the field: NJR, BOB. Taxonomic analysis of samples: NJR, EAL-W, and JDZ. Laboratory analyses: EL-W, EAL-W. Data analysis: NJR.

